# Human leukocyte antigen class II gene diversity tunes antibody repertoires to common pathogens

**DOI:** 10.1101/2021.01.11.426296

**Authors:** Taushif Khan, Mahbuba Rahman, Ikhlak Ahmed, Fatima Al Ali, Puthen Jithesh, Nico Marr

## Abstract

Allelic diversity of HLA class II genes may help maintain humoral immunity against infectious diseases. We investigated the relative contribution of specific HLA class II alleles, haplotypes and genotypes on the variation of antibody responses to a variety of common pathogens in a cohort of 800 adults representing the general Arab population. We found that classical HLA class II gene heterozygosity confers a selective advantage. Moreover, we demonstrated that multiple HLA class II alleles play a synergistic role in shaping the antibody repertoire. Interestingly, associations of HLA-DRB1 genotypes with specific antigens were identified. Our findings suggest that HLA class II gene polymorphisms confer specific humoral immunity against common pathogens, which may have contributed to the genetic diversity of HLA class II loci during hominine evolution.

## Introduction

Originally discovered as the genetic loci responsible for rapid graft rejection, the classical major histocompatibility complex class I (*MHC-I*) and II (*MHC-II*) genes encode glycoproteins responsible for antigen presentation, allowing the immune systems of all jawed vertebrates to discriminate between self and non-self molecules. In humans, the classical *MHC* genes are located with functionally related genes on chromosome region 6p21.3; this cluster of genes is referred to as the human leucocyte antigen (HLA) gene complex (*1*). HLA class I glycoproteins are ubiquitously expressed, contain the functional sites that primarily bind endogenous peptides and contribute to innate immunity by engaging natural killer cell receptors, and to adaptive cellular immunity, through the engagement of the αβ antigen receptors on cytotoxic (CD8^+^) T cells. In contrast, the HLA class II glycoproteins, HLA-DR, -DP and -DQ, are expressed exclusively by antigen presenting cells. These molecules contribute to adaptive immunity by presenting exogenous peptides and engage with the αβ antigen receptors of helper (CD4^+^) T cells, which in turn participate in the activation of naïve B cells (*1*). Thus, the HLA class II glycoproteins play an indirect but critical role in antibody responses to thymus-dependent antigens. Normally, the peptides presented by the HLA class I and II glycoproteins are derived from host proteins that do not elicit any immune responses due to the elimination of self-reactive T cells during their development in the thymus. This process is orchestrated by the interaction of immature T cells with a variety of thymic cell types. However, following infection or in cancer cells, the binding of non-self (pathogen or mutated) peptides by the HLA glycoproteins leads to the activation of naïve or memory T cells (*2, 3*).

In comparison to most other human genes, the classical HLA loci are extremely polymorphic as a consequence of pathogen-driven balancing selection pressure over prolonged time periods. Some of these polymorphisms were shown to precede the speciation of modern humans (*i.e.* trans-species polymorphisms), or were introduced into the human gene pool by admixture between archaic and modern humans (*i.e.* adaptive introgression) (*1, 4, 5*). To date, more than 25,000 HLA allele sequences have been identified (*6*). Variation is highest at sites that define the peptide-binding repertoire (*5*). Multiple selection mechanisms have been proposed to underly this extraordinarily high level of genetic diversity of classical HLA loci, including negative frequency-dependent selection (also referred to as rare allele advantage), heterozygote advantage, and fluctuating selection, none of which are mutually exclusive (*1, 5*). Nevertheless, providing empirical evidence for the underlying selection mechanisms through human studies and evaluating their relative contribution to HLA diversity have not been straightforward (*5*). Similarly, pinpointing causal variant-disease relationships (or causal variant-phenotype relationships) remains a challenge due to the synergistic effects of multiple HLA loci that have related functions, with each of the classical HLA loci on its own exhibiting a high degree of immunological redundancy, as well as due to the density and strong linkage disequilibrium (LD) of HLA genes (*1, 4, 5, 7*).

The functional effects of common polymorphisms in HLA loci or elsewhere in the human genome have mainly been inferred using an epidemiological study design, in which a group of selected cases with a study-defined disease or individuals with a specific immunological phenotype (*e.g.* a vaccine response or lack thereof) are compared to a group of controls to identify those polymorphisms and alleles that are statistically over-represented among either the case group (*i.e.* risk alleles) or the controls (*i.e.* protective alleles). Such studies have revealed associations of certain HLA class I gene polymorphisms with human immunodeficiency virus-type 1 (HIV-1) virus load and AIDS progression (*8, 9*). Associations have also been identified between HLA class II gene variants and chronic hepatitis B and C infections, leprosy and tuberculosis, or responses to influenza and hepatitis B vaccination, albeit most identified risk or protective alleles have only small-to-modest effect sizes. Moreover, specific HLA alleles have been associated with a variety of autoimmune and inflammatory diseases (*10*). These associations highlight the delicate balance between the ability of the immune system to activate potent effector mechanisms against invading pathogens while preventing excessive host tissue damage (*11*).

Nevertheless, our current understanding of the inter-individual variation of the immune responses to microbial challenges remains limited. The relative contribution of different genetic and non-genetic factors driving this variation are only beginning to be unraveled using holistic (*i.e.* systems immunology) approaches applied to larger cohorts of either healthy individuals, or the general population of a given geographic region (or ethnicity). These approaches allow the dissection of gene-phenotype relationships underlying the enormous inter-individual differences in susceptibility to pathogens at a much higher resolution (*12*). To date, only a few studies have investigated the functional consequences of genetic variation in HLA class II genes on the variability of antibody responses in healthy individuals or the general population (*13–15*). Such studies have been hampered not only by the large number of different HLA class II alleles, the strong LD and the high immunological redundancy of individual HLA class II genes, but also the lack of cost-effective and technically feasible experimental approaches that enable the assessment of very large numbers of antibody-antigen interactions in sufficiently sized human cohorts.

In this study, we explored the relative contribution of specific HLA class II alleles, haplotypes and genotypes on the variation of human antibody responses to a variety of common human pathogens. We conducted an unbiased, large-scale, high-throughput screen of antigen-antibody interactions using phage-immunoprecipitation sequencing (PhIP-Seq) (*16, 17*) and samples from a well-defined cohort of 800 adult Qatari nationals and long-term residents of Qatar. This sample of the general population was expected to have limited genetic diversity and an excess of individuals with HLA homozygosity due to high rates of consanguinity (*18*), thereby allowing us to overcome challenges related to the extreme allelic diversity of classical HLA class II loci.

## Results

### HLA type inference from whole genome sequencing data of the 800 study participants

Using a population reference graph (PRG) framework as described by Dilthey *et al*. (*19*), we determined the allelic state of the classical HLA class II genes in our study cohort of 800 Qatari nationals and long-term residents of Qatar based on whole genome sequencing data at 6-digit allelic resolution or higher (*i.e.* taking into account non-synonymous and synonymous single nucleotide variants in the protein-coding region of the classical HLA class II genes). As expected, *HLA-DRB1* was the most polymorphic gene among the HLA class II genes, with 49 different alleles identified in our cohort, followed by *HLA-DPB1* (28 alleles), *HLA-DPA1* (22 alleles) and *HLA-DQB1* (16 alleles). The most commonly present *DRB1* alleles (≥10%) were HLA-DRB1*03:01:01 (15.81%), HLA-DRB1*07:01:01, (15.06%) and HLA-DRB1*16:02:01 (10.50%). Of note, more than half of our study cohort shared one of two HLA-DRB1 alleles (HLA-DRB1*03:01:01 and HLA-DRB1*07:01:01). Interestingly, we also identified several null alleles in HLA-DRB1 heterozygotes of our study cohort, including HLA-DRB1*15:13 (allele frequency (AF) = 3.19%, n = 51) HLA-DRB1*15:96 (AF = 0.88%, n = 14), HLA-DRB1*07:10 (AF = 0.82%, n = 13) and four more rare HLA-DRB1 alleles (AF <0.2%, not shown). Table 1 lists all detected HLA class II alleles with an estimated AF of 0.5% or more in our study cohort. A multiple sequence alignment of the gene products of all HLA-DRB1 alleles analyzed in this study is shown in Supplementary Figure S1. As expected, genetic variants in the class II loci were found to be in strong linkage disequilibrium (LD) (Supplementary Figure S2), *i.e.* the HLA class II alleles are strongly associated in the population and are inherited as haplotypes (Supplementary Table S1). Due to the high rates of consanguineous marriages in Qatar (*20*), we also assessed the existence of a significant deviation in the observed number HLA homozygotes for the classical class II alleles in our study cohort, assuming Hardy–Weinberg equilibrium. Indeed, excess homozygosity was found for DRB3*03:01:01G, DRB3*01:01:02G and DRB3*02:02:01G, which was consistent with the low fixation index (mean F = 0.0114; SD = 0.04), indicating that genetic material has been shared in this population through high levels of inbreeding. Intriguingly, we also found that homozygotes of HLA-DRB1*07:01:01 were significantly underrepresented (*P* < 0.00001) and completely absent in our study cohort, suggesting that this genotype is under negative selective pressure (Supplementary Table S2).

**Table 1.** HLA class II alleles in the Qatar Biobank cohort (n = 800) with an estimated allele frequency ≥0.5%.

| Allele | Gene | n (heterozygotes) | n (homozygotes) | n (total)<sup>A</sup> | AF<sup>B</sup> |
| --- | --- | --- | --- | --- | --- |
| DPA1*01:03:01G | DPA1 | 304 | 405 | 709 | 69.63% |
| DPA1*02:01:01; DPA1*02:01:08 | DPA1 | 194 | 47 | 241 | 18.00% |
| DPA1*02:02:02 | DPA1 | 80 | 3 | 83 | 5.38% |
| DPA1*02:01:02 | DPA1 | 25 | 0 | 25 | 1.56% |
| DPA1*03:03 | DPA1 | 18 | 0 | 18 | 1.13% |
| DPA1*02:01:03 | DPA1 | 18 | 0 | 18 | 1.13% |
| DPA1*02:02:05 | DPA1 | 13 | 0 | 13 | 0.81% |
| DPA1*02:01:06 | DPA1 | 10 | 0 | 10 | 0.63% |
| DPB1*04:01:01G | DPB1 | 351 | 130 | 481 | 38.19% |
| DPB1*02:01:02G | DPB1 | 230 | 30 | 260 | 18.13% |
| DPB1*03:01:01G | DPB1 | 151 | 15 | 166 | 11.31% |
| DPB1*14:01:01 | DPB1 | 114 | 4 | 118 | 7.63% |
| DPB1*13:01:01G | DPB1 | 68 | 9 | 77 | 5.38% |
| DPB1*04:02:01G | DPB1 | 71 | 3 | 74 | 4.81% |
| DPB1*01:01:01G | DPB1 | 53 | 2 | 55 | 3.56% |
| DPB1*17:01:01G | DPB1 | 42 | 0 | 42 | 2.63% |
| DPB1*10:01 | DPB1 | 27 | 5 | 32 | 2.31% |
| DPB1*09:01:01 | DPB1 | 27 | 1 | 28 | 1.81% |
| DPB1*05:01:01G | DPB1 | 15 | 0 | 15 | 0.94% |
| DPB1*15:01:01G | DPB1 | 12 | 0 | 12 | 0.75% |
| DQA1*05:01:01G | DQA1 | 308 | 60 | 368 | 26.75% |
| DQA1*01:02:01G | DQA1 | 275 | 68 | 343 | 25.69% |
| DQA1*02:01 | DQA1 | 214 | 34 | 248 | 17.63% |
| DQA1*03:01:01G | DQA1 | 179 | 29 | 208 | 14.81% |
| DQA1*01:01:01G | DQA1 | 100 | 8 | 108 | 7.25% |
| DQA1*01:03:01G | DQA1 | 83 | 3 | 86 | 5.56% |
| DQA1*04:01:01G | DQA1 | 27 | 0 | 27 | 1.69% |
| DQA1*06:01:01G | DQA1 | 9 | 0 | 9 | 0.56% |
| DQB1*02:01:01G | DQB1 | 355 | 92 | 447 | 33.69% |
| DQB1*05:02:01G | DQB1 | 172 | 37 | 209 | 15.38% |
| DQB1*03:01:01G | DQB1 | 166 | 10 | 176 | 11.63% |
| DQB1*03:02:01G | DQB1 | 153 | 22 | 175 | 12.31% |
| DQB1*05:01:01G | DQB1 | 88 | 6 | 94 | 6.25% |
| DQB1*06:02:01G | DQB1 | 70 | 4 | 74 | 4.88% |
| DQB1*06:03:01G | DQB1 | 58 | 3 | 61 | 4.00% |
| DQB1*06:04:01G | DQB1 | 40 | 3 | 43 | 2.88% |
| DQB1*04:02:01G | DQB1 | 40 | 0 | 40 | 2.50% |
| DQB1*06:01:01G | DQB1 | 39 | 0 | 39 | 2.44% |
| DQB1*05:03:01G | DQB1 | 22 | 0 | 22 | 1.38% |
| DQB1*03:03:02G | DQB1 | 18 | 0 | 18 | 1.13% |
| DQB1*06:09:01G | DQB1 | 15 | 0 | 15 | 0.94% |
| DRB1*07:01:01G | DRB1 | 241 | 0 | 241 | 15.06% |
| DRB1*03:01:01G | DRB1 | 219 | 17 | 236 | 15.81% |
| DRB1*16:02:01 | DRB1 | 144 | 12 | 156 | 10.50% |
| DRB1*15:01:01G | DRB1 | 82 | 4 | 86 | 5.63% |
| DRB1*04:03:01 | DRB1 | 86 | 0 | 86 | 5.38% |
| DRB1*04:02:01 | DRB1 | 67 | 0 | 67 | 4.19% |
| DRB1*13:02:01 | DRB1 | 56 | 4 | 60 | 4.00% |
| DRB1*13:01:01G | DRB1 | 52 | 2 | 54 | 3.50% |
| DRB1*15:13 | DRB1 | 51 | 0 | 51 | 3.19% |
| DRB1*11:01:01G | DRB1 | 45 | 1 | 46 | 2.94% |
| DRB1*11:04:01G | DRB1 | 44 | 1 | 45 | 2.88% |
| DRB1*16:01:01 | DRB1 | 43 | 0 | 43 | 2.69% |
| DRB1*10:01:01 | DRB1 | 41 | 0 | 41 | 2.56% |
| DRB1*01:01:01G | DRB1 | 30 | 2 | 32 | 2.13% |
| DRB1*15:02:01 | DRB1 | 28 | 0 | 28 | 1.75% |
| DRB1*04:05:01 | DRB1 | 27 | 0 | 27 | 1.69% |
| DRB1*03:02:01 | DRB1 | 23 | 1 | 24 | 1.56% |
| DRB1*01:02:01G | DRB1 | 23 | 0 | 23 | 1.44% |
| DRB1*13:03:01G | DRB1 | 18 | 1 | 19 | 1.25% |
| DRB1*15:03:01G | DRB1 | 19 | 0 | 19 | 1.19% |
| DRB1*08:04:01 | DRB1 | 19 | 0 | 19 | 1.19% |
| DRB1*14:01:01G | DRB1 | 14 | 0 | 14 | 0.88% |
| DRB1*15:96 | DRB1 | 14 | 0 | 14 | 0.88% |
| DRB1*11:01:02 | DRB1 | 14 | 0 | 14 | 0.88% |
| DRB1*07:10N | DRB1 | 13 | 0 | 13 | 0.81% |
| DRB1*04:01:01 | DRB1 | 12 | 0 | 12 | 0.75% |
| DRB1*12:01:01G | DRB1 | 11 | 0 | 11 | 0.69% |
| DRB1*04:06:01G | DRB1 | 11 | 0 | 11 | 0.69% |
| DRB1*11:02:01 | DRB1 | 9 | 1 | 10 | 0.69% |
| DRB3*01:01:02G | DRB3 | 32 | 386 | 418 | 50.25% |
| DRB3*02:02:01G | DRB3 | 47 | 308 | 355 | 41.44% |
| DRB3*03:01:01G | DRB3 | 28 | 48 | 76 | 7.75% |
| DRB3*02:01:01G | DRB3 | 4 | 2 | 6 | 0.50% |
| DRB4*03:01N | DRB4 | 282 | 393 | 675 | 66.75% |
| DRB4*01:01:01G | DRB4 | 284 | 105 | 389 | 30.88% |
| DRB4*01:03:03 | DRB4 | 35 | 1 | 36 | 2.31% |

<sup>A</sup>Total allele count in the study cohort (n = 800); <sup>B</sup>Allele frequency

### Characterization of antibody responses to common human pathogens

Next, we performed PhIP-Seq (*16, 17*) on serum samples obtained from each individual (n = 800) of our study cohort at a single time-point (*i.e.*, at the time of recruitment by the Qatar Biobank study (*18*)). In brief, this technology enabled us to obtain a comprehensive profile of antibody repertoires in our study cohort using phage display of oligonucleotide-encoded peptides, followed by immunoprecipitation and massive parallel sequencing (*16, 17*). The VirScan phage library used for PhIP-Seq in the present study comprised peptide tiles of up to 56 amino acids in length that overlap by 28 amino acids and collectively encompass the full proteomes of most known human-tropic viruses (approximately 400 species) plus many bacterial protein antigens (*21*). Using this technique, we identified the antibody repertoires of 798 individuals; data from two individuals were excluded from the downstream analysis as these did not meet our stringent criteria for quality control (*22*). We also excluded antibody specificities to species for which we found the seroprevalence in the local adult population to be below 5% (for details see the Materials and Methods section). We retained antibody specificities against a total of 48 microbial species for our downstream analysis (Table 2). As expected, the majority of individuals were seropositive for antibodies against various human-tropic viruses that frequently cause upper respiratory tract infections (*i.e.* ‘common cold’ viruses), and human herpesvirus (HHV) species, which commonly establish life-long persistent infections (*i.e.* latency), as well as bacteria such as *Staphylococcus aureus, Streptococcus pneumoniae, and Mycoplasma pneumoniae*, which frequently colonize the skin or upper airways but that are typically innocuous. We also detected antibodies against human papillomaviruses (HPVs), which cause common warts, enteroviruses (EV) (*i.e.*, EV-A, -B and -C), rotavirus A and *Helicobacter pylori*, which can cause gastrointestinal disease, as well as antibody responses that are likely to reflect immunity from childhood vaccination (*e.g.* to smallpox and polio vaccine strains) (Table 2).

**Table 2.** Frequently detected anti-microbial antibody responses.

| Species | Overall (%) | Female (%) | Male (%) |
| --- | --- | --- | --- |
| *Streptococcus pneumoniae* | 95.9 | 96.1 | 95.5 |
| Rhinovirus B | 93.7 | 93.5 | 94.1 |
| Human herpesvirus 4 | 93.0 | 93.5 | 92.0 |
| *Staphylococcus aureus* | 92.9 | 93.7 | 91.3 |
| Human herpesvirus 5 | 90.2 | 92.6 | 86.1 |
| Human herpesvirus 1 | 74.1 | 76.1 | 70.4 |
| Rhinovirus A | 73.8 | 70.6 | 79.4 |
| Human respiratory syncytial virus | 68.4 | 68.7 | 67.9 |
| Human adenovirus C | 56.6 | 59.3 | 51.9 |
| *Mycoplasma pneumoniae* | 53.8 | 53.0 | 55.1 |
| Human herpesvirus 6B | 47.6 | 53.0 | 38.0 |
| Human parainfluenza virus 3 | 44.4 | 44.8 | 43.6 |
| Human herpesvirus 3 | 43.0 | 41.5 | 45.6 |
| Human herpesvirus 7 | 43.0 | 44.2 | 40.8 |
| Human herpesvirus 2 | 40.2 | 39.7 | 41.1 |
| Enterovirus B | 38.7 | 37.8 | 40.4 |
| Human herpesvirus 8 | 38.6 | 39.5 | 36.9 |
| Influenza A virus | 37.3 | 35.2 | 41.1 |
| Enterovirus A | 35.3 | 33.3 | 39.0 |
| Human metapneumovirus | 34.7 | 34.2 | 35.5 |
| Enterovirus C | 30.2 | 29.4 | 31.7 |
| Influenza B virus | 29.9 | 26.0 | 36.9 |
| Vaccinia virus | 28.4 | 28.4 | 28.6 |
| Human coronavirus HKU1 | 25.9 | 25.8 | 26.1 |
| Norwalk virus | 25.6 | 29.2 | 19.2 |
| Human herpesvirus 6A | 24.2 | 25.0 | 22.6 |
| Human adenovirus F | 24.2 | 22.9 | 26.5 |
| Human adenovirus D | 23.9 | 23.7 | 24.4 |
| *Helicobacter pylori* | 19.5 | 20.5 | 17.8 |
| Cosavirus A | 15.2 | 15.9 | 13.9 |
| Influenza C virus | 14.5 | 15.5 | 12.9 |
| Hepatitis B virus | 14.4 | 14.3 | 14.6 |
| Rotavirus A | 14.4 | 14.1 | 15.0 |
| Alphapapillomavirus 10 | 14.4 | 17.0 | 9.8 |
| Cowpox virus | 14.2 | 15.1 | 12.5 |
| Adeno-associated dependoparvovirus A | 12.8 | 11.4 | 15.3 |
| Human adenovirus B | 12.7 | 11.9 | 13.9 |
| Human parvovirus B19 | 12.7 | 12.9 | 12.2 |
| Alphapapillomavirus 9 | 12.5 | 12.5 | 12.5 |
| Sapporo virus | 12.3 | 11.5 | 13.6 |
| Human parainfluenza virus 1 | 10.8 | 11.5 | 9.4 |
| Aichivirus A | 10.3 | 9.0 | 12.5 |
| Human coronavirus NL63 | 8.9 | 9.2 | 8.4 |
| Human parainfluenza virus 2 | 7.8 | 8.2 | 7.0 |
| Human adenovirus E | 7.6 | 6.7 | 9.4 |
| Alphapapillomavirus 6 | 7.5 | 8.0 | 6.6 |
| Human adenovirus A | 6.9 | 6.3 | 8.0 |
| Human coronavirus 229E | 5.6 | 4.3 | 8.0 |

### Impact of age and sex on the species-specific antibody responses

Previous studies of the French *Milieu Interieur* cohort showed that age and sex are important non-genetic covariates underlying the inter-individual variability of human antibody responses to common pathogens among healthy individuals (*15*). We therefore included age and sex as covariates in the serological analysis of our cohort. We found the breadth of the antibody repertoire against HHV-5 to be significantly and positively associated with age [−log_10_(*P* value) ≤6.48; β = 1.36; 95% CI: 0.94–1.86], whereas the antibody repertoire breadth against human rhinoviruses (HRV)-A and -B, EV-A, human adenovirus (HAdV)-C, HHV-6B and *S. pneumoniae* correlated negatively with increasing age. We also found the antibody repertoire breadth against influenza B virus (IBV) to be weakly associated with male, but not female sex, whereas the oppositive was the case for the antibody repertoire breadth against HHV-8 and *H. pylori* (Supplementary Table S3).

### Zygosity of classical HLA class II genes affects the antimicrobial antibody repertoire breadth

Our study cohort included a small but sizable proportion of HLA-DRB1 homozygotes (n = 46) (Table 1). HLA diversity among these subjects was more limited at the individual level compared to that of the HLA heterozygotes, because these individuals inherited the same HLA-DRB1 allele (as well as HLA haplotypes with low sequence divergence) from each parent. Consequently, they express fewer molecular variants of the HLA-DP, -DQ, and -DR heterodimers that present peptides to CD4^+^ T cells. We therefore reasoned that HLA-DRB1 homozygotes would also have a lower capacity for generating antibody responses against a broad spectrum of antigens than HLA-DRB1 heterozygotes, at least in response to some pathogens, and independent of the specific HLA alleles and haplotypes they have inherited. To account for the varying number of peptides and potential protein antigens for each microbial species encompassed by the phage display library, we adjusted the species-specific score values by normalizing the counts of significantly enriched, non-homologous peptides (*i.e.* pulled down peptides containing distinct linear B cell epitopes) against the total count of peptides for a given microbial species represented in the phage library, as described previously (*22*). We then used these adjusted species score values as a quantitative measure of the antibody repertoire breadth against each of the common microbial species identified, thereby allowing us to independently assess the effect of HLA-DRB1, -DPA1, -DPB1, -DQA1, -DQB1 zygosity. We found a significant (*P* ≤0.0001) and positive (β ≥0.68) association between a heterozygous HLA-DRB1 genotype and the size of the antibody repertoires against HHV-4, HHV-5, HHV-6B, HRV-A, HRV-B and HAdV-C, as well as *S. aureus* and *S. pneumoniae* (Table 3). To account for the imbalance in the sample size for homozygous versus heterozygous individuals, we confirmed these findings using a bootstrapping method after randomly re-sampling (100 times) the same number of heterozygotes and HLA-DRB1 homozygotes (Supplementary Figure 3).

**Table 3.** Associations between zygosity in HLA class II genes and the antibody repertoire breadth of selected species.

| Explanatory variable<sup>A</sup> | Response variable<sup>B</sup> | $-\log_{10}(P\text{-value})$ | Coefficient of association ($\beta$) | 95% CI |
| --- | --- | --- | --- | --- |
| DRB1 zygosity | *Streptococcus pneumoniae* | 42.5 | 2.01 | 1.73 - 2.30 |
| DRB1 zygosity | *Staphylococcus aureus* | 20.3 | 1.51 | 1.19 - 1.82 |
| DRB1 zygosity | Human herpesvirus 5 | 10.6 | 1.00 | 0.71 - 1.30 |
| DRB1 zygosity | Human herpesvirus 4 | 10.2 | 1.39 | 0.98 - 1.81 |
| DRB1 zygosity | Rhinovirus B | 26.8 | 1.34 | 1.10 - 1.59 |
| DRB1 zygosity | Rhinovirus A | 25.9 | 1.06 | 0.87 - 1.26 |
| DRB1 zygosity | Human adenovirus C | 22.1 | 0.89 | 0.72 - 1.07 |
| DRB1 zygosity | Human herpesvirus 6B | 18.4 | 0.78 | 0.61 - 0.95 |

<sup>A</sup> Zygosity in HLA-DRB1, -DPA1, -DPB1, -DQA1, and -DQB1 were used as explanatory variables. Only results with a significant association ( $P\text{-value} \leq 0.0001$ ) are shown; <sup>B</sup> The adjusted species score values (response variables) served as a measure of the antibody repertoire breadth in the selected species.

### HLA class II allele- and HLA-DQA1~DQB1~DRB1 haplotype-specific effects on the antibody repertoire breadth against common microbial infections

Given the pathogen-driven balancing selection and allelic diversity of classical HLA class II loci, particularly at sites that define the peptide-binding repertoire (*5*), we reasoned that classical HLA class II alleles and haplotypes should have varying effects on the repertoire breadth of antibodies detected in our study cohort, which may also differ depending on the microbial species. These microbial species, for which the variance in human antibody responses between individuals with different HLA class II alleles and haplotypes is greatest, may arguably also play an important role in driving allelic diversity of HLA class II genes in the first place. We tested for associations between specific HLA class II alleles and the antibody repertoire breadth against the common microbial species identified above by using the adjusted species score values as response variables and the HLA-DRB1 (n = 21), -DQB1 (n = 11), -DPB1 (n = 9), -DQA1 (n = 5) and -DPA1 (n = 5) alleles with an AF between 1% and 20% in our study cohort as explanatory variables. HLA-DRA alleles were excluded from the analysis as there are no polymorphisms of this gene in sequences encoding the peptide-binding grooves (*1*). We also excluded rare (AF <1%) as well as more common (AF >20%) HLA class II alleles found in our study cohort (i.e. HLA-DRB3 and -DRB4 alleles) to ensure homoscedasticity (not shown). Again, stringent criteria were applied to test for strong associations (|β| ≥0.68) and to account for multiple comparisons (*P* ≤0.0001). We also defined a new feature for each tested HLA class II allele, namely the anti-microbial response ratio (RR), which was calculated by dividing the number of significant and positive associations of the antibody repertoire breadth to multiple microbial species by the total number of microbial species for which we had identified at least one pairwise association (see the Materials and Methods). Our analysis revealed significant and positive associations of 10 HLA class II alleles with the antibody repertoire breadth against at least one of 11 microbial species and no negative associations were identified. Positive associations were most robust [i.e., −log_10_(*P*-value) ≥10 and/or a RR ≥0.3] for HLA class II alleles DRB1*03:01:01G, DQB1*03:01:01G, DRB1*13:02:01 and DQA1*02:01, which were associated with the breadth of the antibody repertoires against multiple microbial species, such as *S. pneumoniae*, *S. aureus*, HRV-A, HRV-B, HHV-4, HHV-5, HHV-6B, HAdV-C and HRSV (Figure 1A). We also performed a regression analysis of the adjusted species-specific scores by using the HLA-DQA1~DQB1~DRB1 haplotypes with a frequency ≥1% (Supplementary Table S1) as explanatory variables. In this way, we identified significant positive associations between 14 haplotypes and the antibody repertoire breadth against 17 microbial species. With the exception of one haplotype, all positively associated haplotypes were represented by at least one of the HLA-DRB1, -DQB1 or -DQA1 alleles for which we had also independently identified an association with the antibody repertoire breadth. Of note, we observed a synergistic effect of multiple HLA class II alleles, as both the magnitude of RR and the strength of association with the antibody repertoires against individual species was higher in comparison to our previous analysis when each allele was assessed separately. Again, positive associations were most common with the antibody repertoire breadth against *S. pneumoniae*, *S. aureus*, HRV-A, HRV-B, HHV-4, HHV-5, HHV-6B, HAdV-C and HRSV. Moreover, we also identified positive associations with the antibody repertoire breadth against EV-A and -B, IAV, human parainfluenza virus (HPIV)-3, HHV-1, HHV-7 and *M. pneumoniae*, further demonstrating a synergistic effect of different HLA class II alleles (Figure 1B).

**Figure 1.**
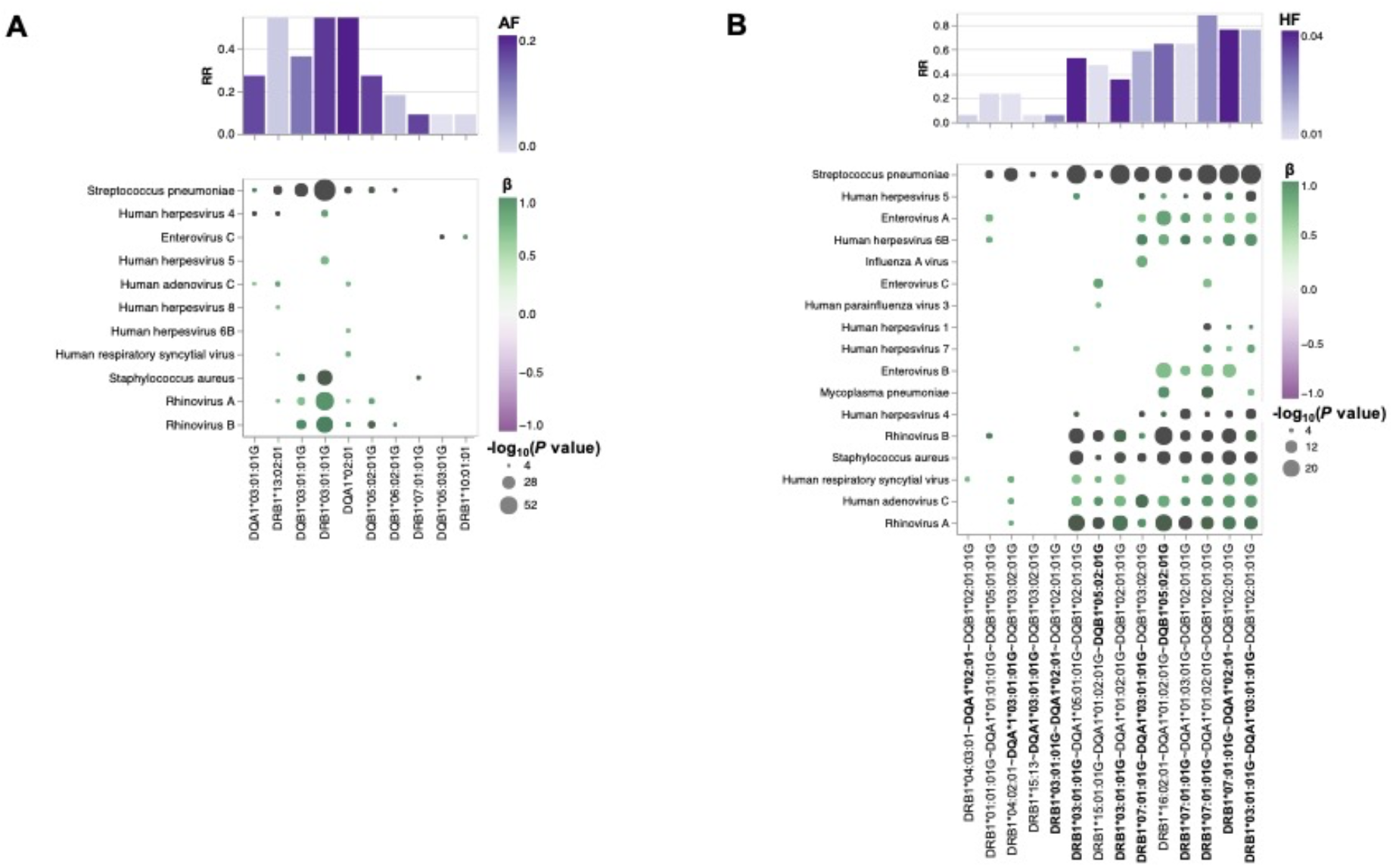
HLA class II allele-specific and HLA-DQA1~DQB1~DRB1 haplotype-specific effects on the antibody repertoire breadth against common microbial infections. **A, B,** Heatmap plots depicting significant associations (*P* < 0.0001) between specific alleles **(A)** or haplotypes **(B)** and the antibody repertoire breadth against common microbial infections. The coefficient (β) and direction of associations are indicated by a color gradient for each circle. The circle size depicts the −log_10_(*P*-value) of the association. Alleles for which significant associations were independently identified as shown in **(A)** are labeled in bold in **(B)**. Bar plots depict the anti-microbial response ratio (RR) for each allele or haplotype. The allele or haplotype frequency is indicated by a color gradient for each bar.

### Associations between HLA-DRB1 genotypes and the antibody repertoire breadth against common microbial infections

Humans are diploid organisms and ultimately, it is likely that the synergistic effect of multiple HLA class II alleles encoded on both parental chromosomes defines the antibody binding repertoire of a given individual with a specific HLA class II genotype and diplotype. However, assessment of the role of all HLA-DQA1~DQB1~DRB1 diplotypes remains a challenge, primarily due to the extremely polymorphic nature of the classical HLA genes. Thus, studies of very large cohorts are required to achieve sufficiently sized groups with identical diplotypes for statistical comparison; this was not feasible in our cohort of 800 individuals. To overcome this issue, we tested for associations between specific HLA-DRB1 genotype groups and the breadth of the antibody repertoires against each of the common microbial species described above, and took advantage of the strong LD of the HLA class II loci (Supplementary Figure S2). We first performed unsupervised clustering of the 800 individuals of our study cohort based on their HLA-DRB1 genotypes using a hierarchical density-based clustering algorithm (HDBSCAN). We focused on *HLA-DRB1*, which is the most polymorphic locus among the HLA class II genes, because it allowed us to assign a maximum number of individuals in our cohort to a specific cluster with significant probability estimation compared to any of the other less polymorphic HLA class II genes (not shown). Approximately 43% of the individuals (n =357) in our study cohort were clustered with significant probability estimation into one of 18 clusters (denoted as HLA-DRB1 genotype groups 1–18) (Supplementary Figure S4A), with most groups representing a subset of closely HLA-matched individuals (Figure 2A) and each group represented by a sample size of ≥12 individuals (Supplementary Figure S4B). HLA-DRB1 genotype groups 14 and 18 exclusively comprised HLA-DRB1*16:02:01 homozygotes (n = 12) and HLA-DRB1*03:01:01G homozygotes (n = 17), respectively; which combined represented approximately 3.7% of our study cohort (Figure 2A and Supplementary Figure S4B). The remaining groups comprised HLA heterozygotes and most groups clustered into one of four supergroups that share a major allele (namely HLA-DRB1*03:01:01G, HLA-DRB1*07:01:01, DRB1*15:01:01G or HLA-DRB1*16:02:01) (Figure 2A). As described previously, individuals homozygous for the HLA-DRB1*07:01:01 allele were completely absent from our study cohort (Table 1 and Supplementary Table S2). HLA-DRB1 groups 1 and 3 included individuals with comparatively higher allelic diversity, representing approximately 3.2% of our study cohort (Figure 2A and Supplementary Figure S4B).

**Figure 2.**
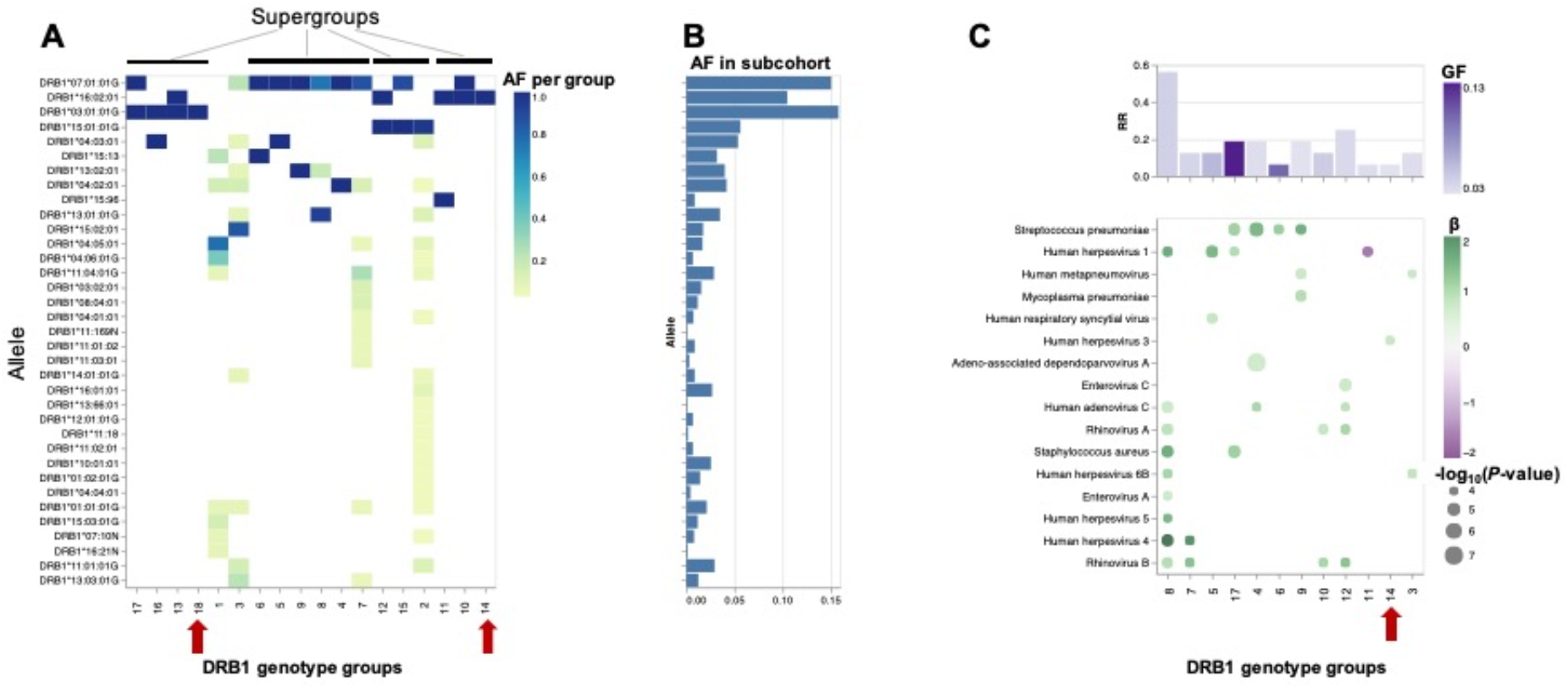
HLA-DRB1 genotype-specific effects on the antibody repertoire breadth against common microbial infections. **A,** Heatmap plot depicting the HLA-DRB1 allele frequency per HLA-DRB1 group. **B,** Bar plot depicting the allele frequency across all individuals assigned to one of eighteen HLA-DRB1 groups (n = 357). **C,** Heatmap plot depicting significant associations (*P* <0.0001) between specific HLA-DRB1 genotype groups and the antibody repertoire breadth against common microbial infections. The coefficient (β) and direction of association is indicated by a color gradient for each circle. The circle size depicts the −log_10_(*P*-value) of the association. Bar plot depicts the anti-microbial response ratio (RR) for each HLA-DRB1 group. The genotype frequency (GF) is indicated by a color gradient for each bar. Groups with HLA-DRB1 homozygotes are indicated by an arrow.

To test for associations between the antibody repertoire breadth against common microbial species and specific *HLA-DRB1* genotypes, we performed a linear regression analysis of the adjusted species score values, this time using the HLA-DRB1 genotype group assignment described above as explanatory variables. Of the 18 HLA-DRB1 genotype groups evaluated, we identified a significant positive association in 11 groups (groups 3–10, 12, 14 and 17) with the antibody repertoire breadth against at least one (RR ≥ 0.0625) and up to nine (RR = 0.5625) out of 16 microbial species. In contrast, heterozygosity for DRB1*16:02:01 and the null allele DRB1*15:96 (group 11) was negatively associated with the antibody repertoire breadth against HHV-1 (Figure 2A and C). Most robust positive associations (RR = 0.0625) were found for individuals in HLA-DRB1 genotype group 8 carrying the DRB1*07:01:01G allele in combination with either DRB1*13:01:01G or DRB1*13:02:01 (Figure 2A and 2C). Notably, no significant associations were found for HLA-DRB1*03:01:01G homozygotes (group 18) and homozygosity for HLA-DRB1*16:02:01 (group 14) was only marginally positively associated with the antibody repertoire breadth against HHV-3 [β = 0.81; −log_10_(*P*-value) = 4.12]. In accordance with the results of our association studies at the allele and haplotype level, positive associations between HLA-DRB1 genotypes and the antibody repertoire breadth were mainly observed for a limited number of microbial species, including bacterial species such as *S. pneumoniae, S. aureus* and *M. pneumoniae,* human herpesviruses (HHV-1, HHV-3, HHV-4, HHV-5, HHV-6B), ‘common cold’ RNA viruses (HRV-A, HRV-B, HRSV, human metapneumovirus [HMPV]) and viruses that also (or primarily) infect the gastrointestinal tract of humans (HAdV-C, EV-A, EV-C), but usually cause only mild or no symptoms.

### Associations of HLA-DRB1 genotypes with specific antigens

Finally, we sought to assess the effect of specific HLA-DRB1 alleles and genotypes at the antigen level. The gene products of advantageous HLA alleles and genotypes may not only be able to present a broader array of pathogen-derived peptides than risk alleles and genotypes, but may also enhance the peptide-binding specificity and presentation of selected antigenic regions (*i.e.* epitopes) (*23*). To explore the effect of HLA-DRB1 genotypes on antibody binding specificities to common microbial antigens, we first filtered for peptide antigens that were significantly enriched in at least two samples of our cohort and were also differentially enriched across the different DRB1 groups described above, by using Fisher’s exact test [−log_10_(*P*-value) ≥2.3] and an OR of ≥2 or ≤-2 as the cut-off. Interestingly, most of the variance among the retained differentially enriched peptide antigens was due to antibodies targeting proteins of a relatively few microbial species, most notably HHV-1, HHV-2, HHV-4, HHV-5, IAV, IBV, and HAdVs A-E (Figure 3A-B). We then filtered for protein antigens for which the differentially enriched peptides showed high variance (above the 75th quartile) across the different DRB1 groups (for details see the Materials and Methods section). Following the application of these stringent filter criteria, 28 protein antigens were retained, representing 13 microbial species, most notably HHVs and HAdVs (Figure 3C). Among these microbial antigens, we found considerable variance in the antibody specificities targeting a variety of HHV proteins, including tegument proteins VP22, UL14, US11, VP 16, the envelope protein US9 and glycoprotein I (gI) of herpes simplex virus 1 (HSV-1, species HHV-1), the envelope glycoprotein D of herpes simplex virus 2 (HSV-2, species HHV-2), a VP26 homolog of varicella-zoster virus (VZV, species HHV-3) and the Epstein–Barr virus nuclear antigen 5 (EBNA-5) as well as the SM protein (species HHV-4) (Figure 3B-G). A multiple sequence alignment of these HHV antigens by Clustal Omega did not reveal linear amino acid sequence similarities (not shown), indicating that these antigens are targeted by antibodies with distinct specificities owing to multiple HLA class II alleles. The VP26 homolog of VZV for example, was frequently targeted by individuals in DRB1 genotype group 14 (Figure 3C and 3F) that comprised homozygotes of HLA-DRB1*16:02:01, a genotype we had also found to be associated with the antibody repertoire breadth to the same virus species (Figure 2C). In contrast, we also found that individuals in some DRB1 groups (e.g. groups 4 and 5) had antibodies that frequently targeted antigenic peptides of different HAdV species. All these peptides showed a high degree of amino acid similarity and resembled a region of an orthologous core protein expressed by the different species (Figure 3B, 3C, 3H-J), suggesting similar antibody specificities that may also cross-react with antigens of the other HAdV species.

**Figure 3.**
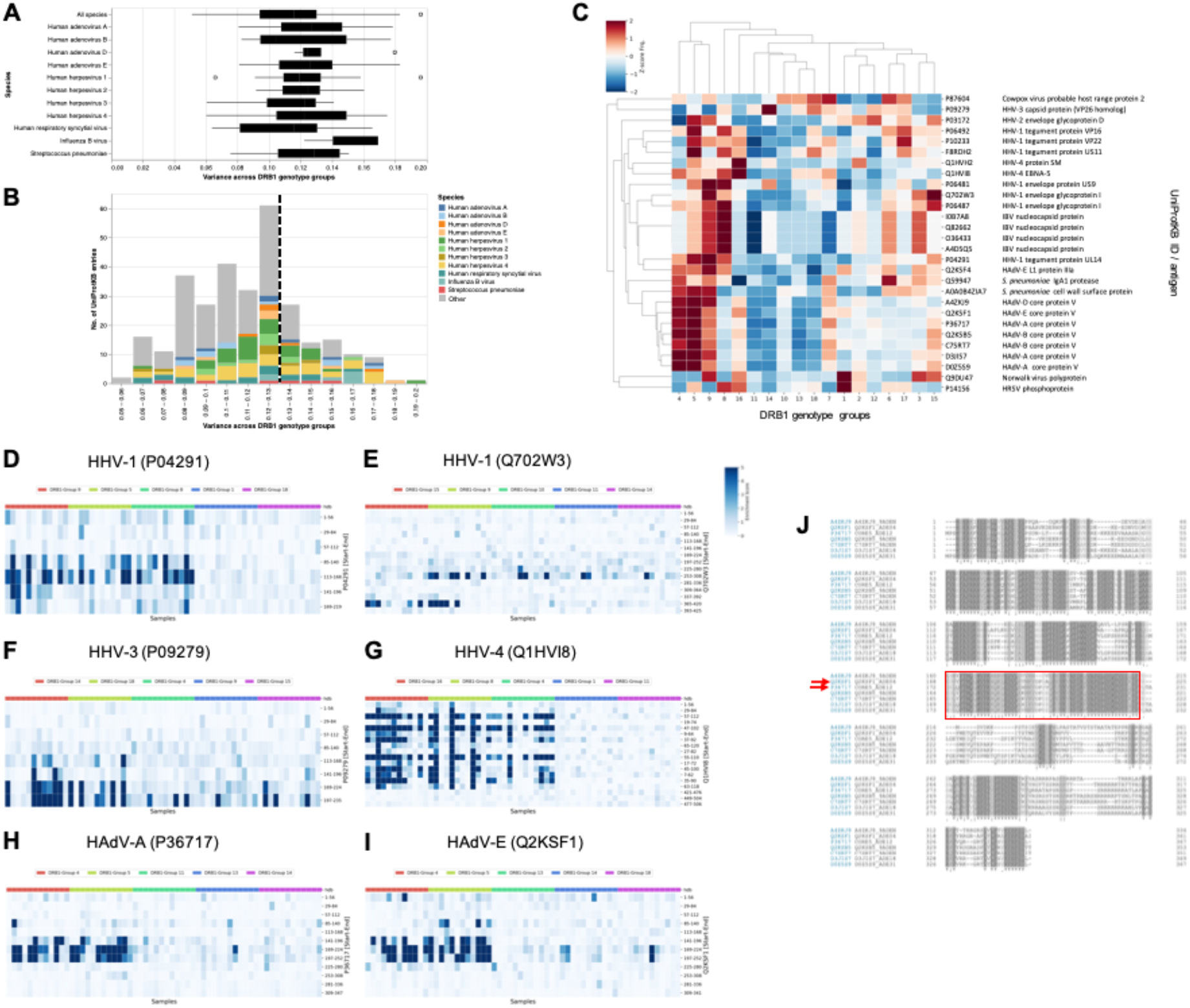
Differential enrichment analysis of antigenic peptide-antibody interactions among different HLA-DRB1 groups. **A,** Boxplot illustrating the distribution of variance in the mean peptide enrichment frequency per UniProtKB entry of all, or selected, microbial species, and across different HLA-DRB1 genotype groups. **B,** Histogram of variance in the mean peptide enrichment frequency per UniProtKB entry, as shown in **(A)**. High variance captures protein antigens of microbial species that exhibit a comparatively different peptide enrichment profile across HLA-DRB1 genotype groups. The dashed line in **(B)** indicates the boundary of the upper 25^th^ percentile (variance ≥0.13). UniProtKB entries for which the antibody-antigen interactions showed the highest variance across different DRB1 groups were color-coded by species. **C,** Heatmap plot showing the antibody binding profile of selected microbial antigens across different HLA-DRB1 groups, with hierarchical clustering. Each row is a protein (UniProtKB entry) with a variance ≥0.13 in the mean peptide enrichment frequency as shown in (**B**); each column represents a HLA-DRB1 genotype group. The color gradient represents the mean enrichment score (Z-score) of antigenic peptides per protein antigen and HLA-DRB1 genotype group. **D-I,** Comparative antigenicity profiles of selected microbial antigens across DRB1 genotype groups. Only representative DRB1 genotype groups with high variance in the Z-score values are shown. In the heatmap plots, each row is a peptide tiling across the indicated protein; each column represents a random sample of the selected DRB1 genotype groups (10 samples per group are shown). Individuals of the same group are indicated with a color bar (top). The color intensity of each cell corresponds to the −log_10_(*P*-value), which was used as a measure of enrichment for a peptide in a sample. Greater values indicate stronger antibody responses; a −log_10_(*P*-value) ≥2.3 was considered to indicate statistical significance. **J**, Multiple sequence alignment (Clustal Omega) of the L2 gene products (Core protein V) of different HAdV species shown in (**C**). Red arrows and the box indicate the UniProtKB entries and antigenic region corresponding to the peptide tiles with strong antibody responses shown in (**H**) and (**I**).

## Discussion

In this study, we employed a systematic and unbiased approach to explore the relative contribution of germline genetic variation in classical HLA class II genes among the general adult population to human antibody responses, including antibody specificities to 48 common human-tropic pathogenic microbial species. By applying a high-throughput method for large-scale antibody profiling to a well-defined cohort of mostly Qatari nationals sharing genetic material as a result of high levels of inbreeding, we dissected the overall effect of zygosity for classical HLA class II genes, as well as effects associated with specific HLA class II alleles, haplotypes and genotypes, on the antimicrobial antibody repertoire breadth and antibody specificity with unprecedented resolution.

Our results provide direct evidence that heterozygosity in classical HLA class II genes confers a selective advantage in humans. Heterozygote advantage has been proposed as one of the main mechanisms that has driven HLA allelic diversity and resistance to infection during human evolution. However, direct empirical evidence from human studies has been sparse (*1, 5*). Our genetic analysis of classical HLA class II allele and genotype frequencies provided the first evidence in support of this mechanism in the context of HLA class II loci. Surprisingly, although HLA-DRB1*07:01:01 was one of the most common DRB1 alleles in our study cohort [AF = 15.06%, n (heterozygotes) = 241], HLA-DRB1*07:01:01 homozygotes were completely absent, suggesting that this genotype is under negative selective pressure (*P* < 0.00001). In contrast, individuals heterozygous for HLA-DRB1*07:01:01 exhibited an antimicrobial antibody profile that was largely indistinguishable from that of individuals who expressed other more or less common DRB1 alleles investigated in this study. When assessed separately, we found HLA-DRB1*07:01:01 was associated only with antibody responses against *S. aureus* (Figure 1A). In contrast, individuals with a haplotype carrying the HLA-DRB1*07:01:01 allele in combination with other HLA-DQA and -DQB alleles (e.g. haplotypes DRB1*07:01:01G~DQA1*03:01:01G~DQB1*03:02:01G or DRB1*07:01:01G~DQA1*01:02:01G~DQB1*02:01:01G) exhibited broad antibody responses that were associated with polyclonal antibody responses to a variety of microbial species (Figure 1B). Similarly, we found a positive association between individuals in DRB1 group 8, with most of them heterozygous for the HLA-DRB1*07:01:01 and DRB1*13:01:01G alleles (Figure 2A), and the antibody repertoire breadth to a variety of microbial species (RR >0.5; Figure 2C). The same individuals also showed strong antibody responses to specific antigens, such as the IBV N protein, the HHV-1 envelope and tegument proteins, the HRSV phosphoprotein, and EBNA-5 (Figure 3C). Taken together, these findings suggest a highly redundant role of the DRB1*07:01 allele in the antibody responses in heterozygous individuals and a compensatory effect of other HLA class II alleles, although this remains to be verified. Of note, the HLA-DRB1*07:01 allele has previously been associated with persistent HCV infection (*24*), as well as asparaginase hypersensitivity and anti-asparaginase antibodies and may therefore lead to suboptimal drug responses and a greater risk of relapse in heterozygous carriers who develop leukemia and lymphomas (*25*). It remains to be determined whether homozygotes for this allele are more prone to certain infectious, allergic or autoimmune diseases, or if this genotype is perhaps associated with other diseases, early death or infertility. The DRB1*07:01 AF in our study cohort is largely comparable or even lower than that reported for other Arab populations and ethnicities, such as Saudis (26.6%), Yemenite-Jews (22.1%), Libyans (17.0%) or Algerians (15.9%) (*26*), suggesting that homozygosity of this allele may represent a common and important genetic risk factor among Arab populations, particularly for children of consanguineous parents.

We also demonstrate that overall (*i.e.*, irrespective of the DRB1 allele), HLA-DRB1 heterozygotes have a broader antibody repertoire against a variety of viral and opportunistic bacterial pathogens, including HHV-4, HHV-5, HHV-6B, HRV-A, HRV-B and HAdV-C, *S. aureus* and *S. pneumoniae,* when compared to HLA-DRB1 homozygotes, which we found to represent a smaller but still sizable proportion (5.75%, n = 46) of our study cohort (Table 1). The relatively high proportion of HLA homozygotes in our study cohort can be explained by the high rates of consanguineous marriage in the State of Qatar (*20*). Finally, we provide evidence of a heterozygote advantage of classical HLA class II loci by a comparative analysis of groups of closely HLA-matched individuals assigned to distinct groups based on their HLA-DRB1 genotypes. Two of these groups comprised HLA-DRB1*16:02:01 homozygotes (group 14, n = 12) or HLA-DRB1*03:01:01G homozygotes (group 18, n = 17) exclusively. Neither of the two groups of HLA-DRB1 homozygotes exhibited antibody responses that were associated with the antibody repertoire breadth or strong antibody responses to specific antigens of multiple microbial species. In contrast, heterozygotes in group 17 expressing the common DRB1*03:01:01G and DRB1*07:01:01G alleles for example, exhibited antibody responses that were positively associated with the antibody repertoire breadth against *S. pneumoniae*, *S. aureus* and HHV-1 (Figure 2), and had stronger antibody responses to specific antigens, such as HHV-1 tegument proteins and a cell wall surface protein of *S. pneumoniae* (Figure 3C). The only significant association we could identify between a homozygous HLA-DRB1*16:02:01 genotype and HHV-3 (Figure 2) was attributable mainly to specific responses against a single viral antigen, namely a small capsomere-interacting protein of VZV and homolog of HSV-1 VP26 (Figure 3C and 3F). To the best of our knowledge, this is the first study to provide empirical evidence of a heterozygote advantage of classical HLA class II genes in humans. Thus far, heterozygote advantage in HLA loci has only been documented in the context of HIV infection, as this virus produces escape variants during chronic infection at a considerable frequency (*1*). Maximum HLA heterozygosity of the classical HLA class I genes *HLA-A*, *-B* and *-C* has been associated with delayed disease onset among HIV-1 infected patients, whereas individuals who were homozygous for one or more loci progressed rapidly to AIDS and death (*27*). Other well-known examples of heterozygote advantage include the recessive disease-causing variants underlying sickle-cell anemia, with one copy of the HbS allele shown to protect heterozygotes from severe forms of malaria (*28*). Interestingly, an *in silico* analysis by Sellis *et al*. (*29*) suggested that a substantial proportion of host adaptive mutations that occur(ed) during human and vertebrate evolution could confer a heterozygote advantage, as rapidly changing environments and genetic variation produce a diversity advantage in diploid organisms that allows them to remain better adapted compared with haploids, despite the fitness disadvantage associated with the occurrence of rare homozygotes (*29*).

Our findings also demonstrate that multiple alleles of the classical HLA class II genes (i.e. HLA-DRB1, -DQA1 and -DQB1) play a synergistic role in shaping the antibody repertoire against microbial pathogens. Indeed, when analyzing each allele in isolation, we found only a limited number of associations between a given allele and the antibody repertoire breadth to a specific microbial species. However, when considering HLA-DQA1~DQB1~DRB1 haplotype-specific responses, we identified additional associations between certain allele combinations and the antimicrobial antibody responses, with most groups of individuals sharing the same haplotype also mounting robust antibody responses to a larger number of microbial species. Our results therefore support the concept that viral infections, along with other infectious diseases, have helped to maintain strong immunity and resistance to common infections during human evolution by promoting diversity in HLA class II alleles and consequently, in B cell-mediated antibody responses (*30*). The reasons why HLA diversity at the individual (host) level remains relatively low have been debated since expression of even more HLA molecules or molecular variants by a given individuum, which may arise through gene duplication events that have occurred throughout vertebrate evolution, would theoretically allow the binding and presentation of even a broader spectrum of antigens, thereby enhancing immunity to infections (of note, this may be the case for some individuals with haplotypes that express additional functional DRB genes, which were not present in our study cohort) (*5*). The associated trade-off effects appear to be the most plausible explanation. Indeed, certain HLA alleles have been shown to play a protective role in the context of certain infectious diseases, while at the same time being associated with an increased risk for autoimmune diseases (*5, 10, 31*). In this regard, it should be noted that the HLA-DRB1*03:01 allele, which was relatively common in our study cohort (AF 15.81%), has been reported to be risk allele for autoimmune hepatitis (AIH) (*32*). AIH may develop not only after hepatitis A, B or C infections, but also following more common infections with HSV-1, EBV, or measles virus. The prevalence of AIH in the general adult population in this study remains unknown.

Interestingly, using our unbiased, large-scale screen and in-depth analysis of antibody specificities to 48 microbial species, we predominantly and repeatedly identified positive associations with antibody responses against members of the *Herpesviridae* family [such as HSV-1 (HHV-1), VZV (HHV-3), EBV (HHV-4), CMV (HHV-5), and roseolavirus (HHV-6B)], *Picornaviridae* (including HRV-A and -B, EV-A, -B and -C), *Paramyxoviridae* (e.g. HRSV, and HMPV), *Adenoviridae* (HAdV-C) and also against opportunistic bacterial pathogens that frequently colonize the upper airways of humans but are typically innocuous (*e.g. S. aureus*, *S. pneumoniae* and *M. pneumoniae*). This raises the question of whether these microbial species have also played a critical role during hominine evolution by driving genetic diversity in the classical HLA class II loci. Recent advances in microbial genetics enabling molecular clock analyses suggest that, although phylogenetically diverse, many if not all of these species have evolved in very close association with their human host, some of them (*e.g.* HSV-1) for millennia; similar findings were obtained for their counterparts infecting primates or other vertebrates (*30*). Indeed, although cross-species transmissions in the more recent past have occurred, it is becoming increasingly evident that most human pathogens have their origins long before the Neolithic era (*33*). A commonly stated hypothesis is that pandemic outbreaks of major human infectious diseases (*e.g.* influenza, hepatitis, tuberculosis, malaria, leishmaniasis, and schistosomiasis) that occurred in the more recent (*i.e.* the post-Neolithic) past, causing considerable morbidity and mortality, have been major driving forces of HLA genetic diversity. While this may be true based on the identification of several positive and negative HLA/MHC associations with these diseases (*10*), the role of other human infectious agents, particularly those that have co-evolved with their human host for much longer periods, should not be neglected simply on the basis that they cause no, or only mild, clinical disease in most cases of (modern) human infection. Even herpesviruses such as HSV-1, EBV or CMV, which are most commonly acquired early in life or during childhood, can cause fatal disease in rare patients, either following primary infection of genetically susceptible individuals (*34*), or reactivated infections in patients with cancer, autoimmune diseases or other comorbidities (*35*). Moreover, infections can have more subtle effects on human reproductive fitness. The effects of these ‘modern human pathogens’ on our hominine ancestors and phylogenetically closest relatives (*i.e.*, archaic humans, such as Neanderthals and Denisovans) that are extinct today are also unknown.

It is also important to highlight the limitations of our study. With our large-scale antibody screening approach, we were primarily able to assess antibody specificities and repertoires to linear epitopes of protein antigens, predominantly of human-tropic viruses. Although there is evidence these include neutralizing and non-neutralizing antibodies (*21*), further investigations are required to elucidate the extent to which these genetic and associated immune phenotypic differences affect clinical outcomes of infection, either by long-term longitudinal studies of even larger human cohorts, or a case-control study of selected diseases.

## Materials and Methods

### Study cohort

The study cohort of 800 adult male and female Qatari nationals and long-term residents of Qatar were randomly selected from a larger cohort of individuals taking part in a longitudinal study of the Qatar Biobank (QBB) (*18*) as described previously (*22*). Relevant demographic data of the study subjects have been described previously (*22*).

### HLA type interference from whole genome sequencing data

Whole genome sequencing (WGS) of our study cohort was performed as part of the Qatar Genome Programme (QGP) (https://qatargenome.org.qa/). Sequencing read data were generated and processed as described elsewhere (*36*). In brief, sequencing libraries were generated from whole blood-derived fragmented DNA using the TruSeq DNA Nano kit (Illumina, Inc., San Diego, USA) and sequence reads were generated using a HiSeq X Ten1 system (Illumina, Inc., San Diego, USA). Primary sequencing data were demultiplexed using bcl2fastq (Illumina) and quality control of the raw data was performed using FastQC [v0.11.2] (Babraham Bioinformatics, Babraham Institute, Cambridge, UK). Sequence reads were aligned to the human reference genome sequence [build GRCh38] using Sentieon Genomics pipeline tools (Sentieon, Inc, San Jose, USA) and HLA type interference was performed using the PRG framework described by Dilthey *et al.* (*19*).

### Genetic fixation index and linkage analysis, population differentiation and homozygosity estimation

The genetic fixation index was calculated using PLINK [v 1.9] (*37*). Linkage disequilibrium (LD) was quantified using eLD (*38*). The expected number of homozygotes for a given HLA class II allele was estimated based on the imputed allele frequencies using PRG and assuming Hardy–Weinberg equilibrium. Deviation from the Hardy–Weinberg equilibrium was assessed using Fisher’s exact test and the Bonferroni method was used to correct for multiple testing. A −log_10_(*P*-value) ≥4.7 was considered to indicate statistical significance.

### Hierarchical density-based clustering by HLA-DRB1 genotypes

A hierarchical density-based clustering algorithm (HDBSCAN) (*39*) was used to assign individuals in our cohort to specific clusters (denoted as HLA-DRB1 genotype groups) with significant probability estimation. In brief, we treated each allele as a feature dimension and generated a hyper-dimensional feature space for each variant found in *HLA-DRB1*. The t-distribution stochastic neighbor embedding (tSNE) method was used for two-dimensional (2D) non-linear projection of the multi-dimensional allele feature space. By combining non-linear dimensionality reduction and density-based unsupervised hierarchical clustering, we identified 18 groups of individuals with similar/matching HLA-DRB1 genotypes that could be clearly distinguished from other clusters; each group had a minimum sample size of 12 (Supplementary Figure S4B). A probability score of ≥0.9 was used as cut-off for the cluster assignment; individuals that could not be assigned to any cluster with significant probability estimation (n = 443) were removed from the downstream analysis.

### Phage immunoprecipitation-sequencing (PhIP-Seq) and peptide enrichment analysis

The VirScan phage library used for PhIP-Seq in the present study had been obtained from S. Elledge (Brigham and Women’s Hospital and Harvard University Medical School, Boston, MA, USA). PhIP-Seq of serum samples from the 800 study subjects and peptide enrichment analysis were performed as described previously (*16, 17, 22*). In brief, we utilized an expanded version (*21*) of the original VirScan phage library described by Xu *et. al. (17)*. Custom sequencing libraries were prepared as previously described (*16*) and sequencing was performed using a NextSeq system (Illumina). To filter for significantly enriched peptides, we imputed −log_10_(*P*-values) by fitting a zero-inflated generalized Poisson model to the distribution of output counts and regressed the parameters for each peptide sequence based on the input read count. Peptides that passed a reproducibility threshold of 2.3 [−log_10_(*P*-value)] in two technical sample replicates were considered significantly enriched. We then computed virus score values as described by Xu *et al.* (*17*) and the scores were finally adjusted by dividing them according to previously established species-specific significance cut-off values (*22*). Samples with an adjusted species score ≥1 were considered seropositive for the corresponding microbial species. The prevalence for each species was calculated as the number of seropositive samples divided by total number of samples in the cohort. Similarly, we estimated seroprevalence values for each sex separately (Table 2). We excluded antibody specificities to species from our downstream analysis for which we have found the seroprevalence in the local adult population to be below 5%.

### Association studies

We examined the contribution of the genetic variation in the classical HLA class II loci to the diversity of the antibody repertoire at different resolutions (*i.e.*, by independently assessing the effect of zygosity, haplotypes, alleles and HLA-DRB1 genotype groups). The adjusted species score values were used as response variables (these values served as a measure of the antibody repertoire breadth against each of the 48 microbial species evaluated in this study), and generalized linear models (GLM) were applied (for details see the supplementary materials). We corrected for multiple testing using the Bonferroni method. Coefficients of association (β) were reported using a natural log scale; a |β| ≥0.68 and a *P*-value ≤0.0001 was considered to indicate statistical significance. We defined the anti-microbial RR as a new feature for the assessment of HLA class II alleles, haplotypes, or HLA-DRB1 genotype groups. The RR for a given HLA class II allele was calculated by dividing the number of significant associations of the allele examined by the total number of microbial species for which we identified at least one significant association to any HLA class II alleles assessed in this study. The RR for haplotypes or HLA-DRB1 genotype groups were calculated similarly.

### Differential enrichment analysis of antibody-antigen interactions across DRB1 genotype groups

To examine the differential enrichment at the peptide and antigen level, we first performed pairwise differential enrichment tests per peptide, accounting for all possible pairwise comparisons of the DRB1 genotype groups identified (n = 18). We considered only peptides that were significantly enriched in at least two samples among the total number of samples tested. Accordingly, for each peptide assessed, we performed 153 pair-wise differential enrichment tests ((n×(n-1))/2). Using these filter criteria, we tested, on average, 3,989 (±150) peptides per DRB1 group-pair and a total of 9,155 enriched peptides when considering all DRB1 genotype groups combined. Next, we tested for differential enrichment of antibody-antigen interactions in each tested DRB1 group-pair using an |OR| ≥2 and a *P*-value ≤0.005 (Fisher’s exact test) as the cut-off. After removing peptides from microbial species with a seroprevalence of less than 5%, 502 differentially enriched peptides (DEP) were used in our downstream analysis. We then assessed the variance of significant antibody-antigen interactions (*i.e.*, per UniProtKB entry) across DRB1 genotype groups. To do so, we first estimated the peptide enrichment frequency of each DEP (n = 502) per DRB1 genotype group. This peptide enrichment frequency was calculated as the ratio of the number of samples in the DRB1 genotype group for which a DEP was significantly enriched, divided by the total number of samples in that group. Next, we calculated the mean of the peptide enrichment frequency per UniProtKB entry for each DRB1 group. Finally, we assessed the variance in this mean value for each Uniprot entry and DRB1 group to identify the antibody-antigen interactions with the highest variance across different DRB1 groups. For this purpose, we only considered UniProtKB entries for which the variance distribution was above the 75th quartile and at least two DEP were identified. Finally, we filtered for UniProtKB entries for which DEPs were less frequent (<5 %) among individuals in least one of the DRB1 groups.

### Study approval

The human subject research described here was approved by the institutional research ethics boards of Sidra Medicine and the Qatar Biobank. This included the receipt of written informed consent from all study participants at the recruitment site (Qatar Biobank).

### Data and code availability

All processed data are available in the manuscript or the supplementary materials. Raw reads from PhIP-Seq are made available in NCBI’s Sequence Read Archive (Accession: PRJNA685111) upon publication of the paper. Python in-house scripts used in this study are available upon request. The pipeline for processing the PhIP-Seq data has been published previously (*16*). Raw WGS data of the study participants are accessible through the Qatar Genome Programme (https://qatargenome.org.qa).

## Author contribution statement

NM conceived the study and supervised the project. MR and FA designed and performed experiments. TK developed the data analysis tools for the association studies and differential enrichment analysis. TK, NM and IA analyzed and interpreted the data. PJ co-supervised the HLA variant analysis. NM and TK wrote the paper. All authors have seen and approved the manuscript, which has not been accepted or published elsewhere.

## Acknowledgments

We thank the QBB study participants who provided samples and data for this study. We also thank Qatar Genome and the QBB management and staff, in particular Nahla Afifi, Said I. Ismail and Elizabeth Jose, for allowing us to access and analyze QBB/QGP samples and data; the Integrated Genomics Services team of Sidra Genomics for generating and processing WGS data of study participants; Stephen Elledge (Brigham and Women’s Hospital, Harvard University Medical School) for kindly providing the VirScan phage library used in this study and for his early discussions related to this work; Tomasz Kula (Brigham and Women’s Hospital, Harvard University Medical School) and Benjamin Larman (Johns Hopkins School of Medicine) for their advice on technical aspects related to the PhIP-Seq experiments, as well as Jessica Tamanini (Insight Editing London) for proofreading and editing the manuscript. This work was supported by a grant from the Qatar National Research Fund (grant no. PPM1-1220-150017).

