## Supplemental Text, Tables S1 to S3 and Supplemental Figures S1 to S4 for "Human leukocyte antigen class II gene diversity tunes antibody repertoires to common pathogens"

December 17, 2020

**This PDF file includes:**

Supplementary Text

Tables S1 to S3

Figures S1 to S4

### Supplementary Methods:

#### *Associations with non-genetic features*

To test for associations between the antibody repertoire breadth and age as well as sex, we used a generalized linear model (GLM) with the following equation:

$$\text{Adjusted species score} = \beta_1 * variable + \beta_{age} * Age_{norm} + \beta_{sex} * [sex]$$

Here, the adjusted species score values were used as dependent variables. Age and sex were used as independent variables. We assessed associations for each species in independent tests and corrected for multiple testing using the Bonferroni method. A coefficient of association (|β|) ≥0.68 and a *P*-value ≤0.0001 was considered to indicate statistical significance.

#### *Associations with zygosity*

To test for associations between the antibody repertoire breadth and zygosity of HLA class II genes, we used following equation:

$$\text{Adjusted species score} = \beta_1 * variable + \sum_{i}^{HLA-Class\ II\ genes} \beta_i * gene_i + \beta_3 * covariates + \epsilon_1$$

As described above, we used the adjusted species score values as dependent variables. Zygosity in HLA class II genes DPA1, DPB1, DQA1, DQB1, DRB1 were used as independent variables. Age was used as a covariate. As detailed above, a coefficient of association (|β|) ≥0.68 and a *P*-value ≤0.0001 was considered to indicate statistical significance. We further performed bootstrapping (100-fold) to control for the imbalance in the sample distribution between in DRB1 gene heterozygous and homozygous individuals. For this purpose, we randomly re-sampled the same number of heterozygotes as there were homozygotes (n = 46), and applied the same GLM model as detailed above.

#### *Associations with HLA class II alleles and haplotypes*

To test for associations between the antibody repertoire breadth and HLA class II alleles or HLA-DQA1~DQB1~DRB1 haplotypes, we used the following equations:

$$\text{Adjusted species score} = \beta_1 * variable + \sum_{i}^{haplotype} \beta_i * haplotype_i + \beta_3 * covariates + \epsilon_1$$

$$\text{Adjusted species score} = \beta_1 * variable + \sum_{i}^{allele} \beta_i * allele_i + \beta_3 * covariates + \epsilon_1$$

Only HLA class II alleles and HLA-DQA1~DQB1~DRB1 haplotypes with a frequency ≥1% were assessed. Again, a coefficient of association (|β|) ≥0.68 and a *P*-value ≤0.0001 was considered to indicate statistical significance.

#### *Associations with HLA class II alleles and DRB1 genotypes*

To test for associations between the antibody repertoire breadth and DRB1 genotypes, we used the same approach and formula as described for HLA class II alleles or haplotypes.

### Supplementary Tables

**Supplementary Table S1. HLA-DRB1~DQA1~DQB1 haplotypes in the study cohort (n = 800) with an estimated haplotype frequency ≥1%**

| No. | HLA-DRB1~DQA1~DQB1 haplotype | | | n (total)[A] | HF[B] |
|---|---|---|---|---|---|
| **1** | DRB1*07:01:01G | DQA1*02:01 | DQB1*02:01:01G | 70 | 0.044 |
| **2** | DRB1*03:01:01G | DQA1*05:01:01G | DQB1*02:01:01G | 69 | 0.043 |
| **3** | DRB1*03:01:01G | DQA1*01:02:01G | DQB1*02:01:01G | 68 | 0.043 |
| **4** | DRB1*16:02:01 | DQA1*01:02:01G | DQB1*05:02:01G | 54 | 0.034 |
| **5** | DRB1*16:02:01 | DQA1*05:01:01G | DQB1*05:02:01G | 49 | 0.031 |
| **6** | DRB1*07:01:01G | DQA1*01:02:01G | DQB1*02:01:01G | 44 | 0.028 |
| **7** | DRB1*03:01:01G | DQA1*02:01 | DQB1*02:01:01G | 43 | 0.027 |
| **8** | DRB1*07:01:01G | DQA1*05:01:01G | DQB1*02:01:0*1G | 39 | 0.024 |
| **9** | DRB1*03:01:01G | DQA1*03:01:01G | DQB1*02:01:01G | 34 | 0.021 |
| **10** | DRB1*07:01:01G | DQA1*03:01:01G | DQB1*03:02:01G | 34 | 0.021 |
| **11** | DRB1*15:13 | DQA*01:02:01 | DQB1*02:01:01G | 31 | 0.019 |
| **12** | DRB1*04:03:01 | DQA*01:02:01 | DQB1*02:01:01G | 23 | 0.014 |
| **13** | DRB1*11:01:01G | DQA1*05:01:01G | DQB1*03:01:01G | 22 | 0.014 |
| **14** | DRB1*11:04:01G | DQA1*05:01:01G | DQB1*03:01:01G | 20 | 0.013 |
| **15** | DRB1*01:01:01G | DQA1*01:01:01G | DQB1*05:01:01G | 19 | 0.012 |
| **16** | DRB1*07:01:01G | DQA1*01:03:01G | DQB1*02:01:01G | 18 | 0.011 |
| **17** | DRB1*15:01:01G | DQA1*01:02:01G | DQB1*05:02:01G | 17 | 0.011 |
| **18** | DRB1*04:02:01 | DQA*1*03:01:01G | DQB1*03:02:01G | 16 | 0.010 |
| **19** | DRB1*15:13 | DQA1*03:01:01G | DQB1*03:02:01G | 16 | 0.010 |
| **20** | DRB1*16:02:01 | DQA1*02:01 | DQB1*05:02:01G | 16 | 0.010 |

[A] Total haplotype count in the study cohort (n = 800); [B] Haplotype frequency

**Supplementary Table S2. Excess and under-represented alleles in the study cohort (n = 800)**

| Allele | AF | n (heterozygotes) observed | n (homozygotes) observed | n (homozygotes) expected[A] | $-\log_{10}(P\text{-value})^{B}$ |
|---|---|---|---|---|---|
| DRB1*07:01:01G | 0.15 | 241 | 0 | 18 | 5.16 |
| DRB3*03:01:01G | 0.08 | 28 | 48 | 5 | 10.15 |
| DRB3*01:01:02G | 0.50 | 32 | 386 | 202 | 20.88 |
| DRB3*02:02:01G | 0.41 | 47 | 308 | 137 | 21.00 |

[A] Number of expected homozygotes assuming Hardy–Weinberg equilibrium; [B] *P*-values using Fisher's exact test. A $-\log_{10}(P\text{-value}) \geq 4.7$ was considered to indicate statistical significance after correcting for multiple testing using the Bonferroni method. AF, allele frequency.

**Supplementary Table S3. Results of a multivariate association analysis of age and sex with the antibody repertoire breadth against selected species (n = 48).**

| Feature[A] | Species[B] | Coefficient of association (beta) | 95% CI min. | 95% CI max. |
|---|---|---|---|---|
| Age | Human herpesvirus 5 | 1.41 | 0.93 | 1.89 |
| Age | Human herpesvirus 4 | 1.32 | 0.65 | 2.00 |
| Age | Human herpesvirus 1 | 1.20 | 0.68 | 1.72 |
| Age | Human herpesvirus 2 | 1.10 | 0.54 | 1.66 |
| Age | Enterovirus A | -0.75 | -0.98 | -0.52 |
| Age | Human herpesvirus 6B | -0.76 | -1.03 | -0.49 |
| Age | Rhinovirus B | -0.91 | -1.27 | -0.55 |
| Age | Human adenovirus C | -0.95 | -1.22 | -0.68 |
| Age | Rhinovirus A | -1.10 | -1.39 | -0.81 |
| Age | Streptococcus pneumoniae | -1.36 | -1.77 | -0.95 |
| Female | Streptococcus pneumoniae | 3.33 | 3.10 | 3.55 |
| Female | Staphylococcus aureus | 2.68 | 2.42 | 2.94 |
| Female | Human herpesvirus 4 | 2.46 | 2.10 | 2.83 |
| Female | Rhinovirus B | 2.28 | 2.09 | 2.47 |
| Female | Human herpesvirus 5 | 1.83 | 1.57 | 2.09 |
| Female | Rhinovirus A | 1.81 | 1.66 | 1.97 |
| Female | Human adenovirus C | 1.62 | 1.47 | 1.77 |
| Female | Human herpesvirus 6B | 1.39 | 1.25 | 1.54 |
| Female | Human respiratory syncytial virus | 1.24 | 1.09 | 1.39 |
| Female | Human herpesvirus 1 | 1.18 | 0.90 | 1.46 |
| Female | Mycoplasma pneumoniae | 1.09 | 0.91 | 1.28 |
| Female | Enterovirus B | 1.08 | 0.98 | 1.18 |
| Female | Human herpesvirus 7 | 1.05 | 0.85 | 1.25 |
| Female | Enterovirus A | 0.98 | 0.86 | 1.10 |
| Female | Enterovirus C | 0.93 | 0.80 | 1.05 |
| Female | Human herpesvirus 3 | 0.80 | 0.68 | 0.93 |
| Female | Human parainfluenza virus 3 | 0.74 | 0.58 | 0.91 |
| Female | Influenza A virus | 0.74 | 0.62 | 0.86 |
| Female | Helicobacter pylori (Campylobacter pylori) | 0.68 | 0.48 | 0.89 |
| Male | Streptococcus pneumoniae | 3.13 | 2.89 | 3.37 |
| Male | Staphylococcus aureus | 2.61 | 2.34 | 2.89 |
| Male | Rhinovirus B | 2.47 | 2.27 | 2.68 |
| Male | Human herpesvirus 4 | 2.19 | 1.80 | 2.57 |
| Male | Rhinovirus A | 2.03 | 1.87 | 2.20 |
| Male | Human herpesvirus 5 | 1.66 | 1.38 | 1.93 |

| | | | | |
|---|---|---|---|---|
| Male | Human adenovirus C | 1.49 | 1.34 | 1.65 |
| Male | Human herpesvirus 1 | 1.31 | 1.01 | 1.61 |
| Male | Human respiratory syncytial virus | 1.23 | 1.07 | 1.39 |
| Male | Human herpesvirus 6B | 1.21 | 1.06 | 1.37 |
| Male | Mycoplasma pneumoniae | 1.15 | 0.96 | 1.35 |
| Male | Enterovirus B | 1.13 | 1.02 | 1.23 |
| Male | Human herpesvirus 7 | 1.07 | 0.86 | 1.29 |
| Male | Enterovirus A | 1.05 | 0.92 | 1.18 |
| Male | Enterovirus C | 1.03 | 0.90 | 1.17 |
| Male | Human herpesvirus 3 | 0.83 | 0.70 | 0.96 |
| Male | Influenza A virus | 0.81 | 0.68 | 0.93 |
| Male | Human parainfluenza virus 3 | 0.72 | 0.55 | 0.89 |
| Male | Hepatitis C virus | 0.70 | 0.47 | 0.93 |
| Male | Influenza B virus | 0.68 | 0.57 | 0.80 |

[A] Normalized age, male and female sex were used as explanatory variables. Only results with a significant association (*P*-value ≤0.0001) are shown; [B] Adjusted species score values (response variables) were used as a measure of antibody repertoire breadth against the listed species**.**

### Supplementary Figures

**A**

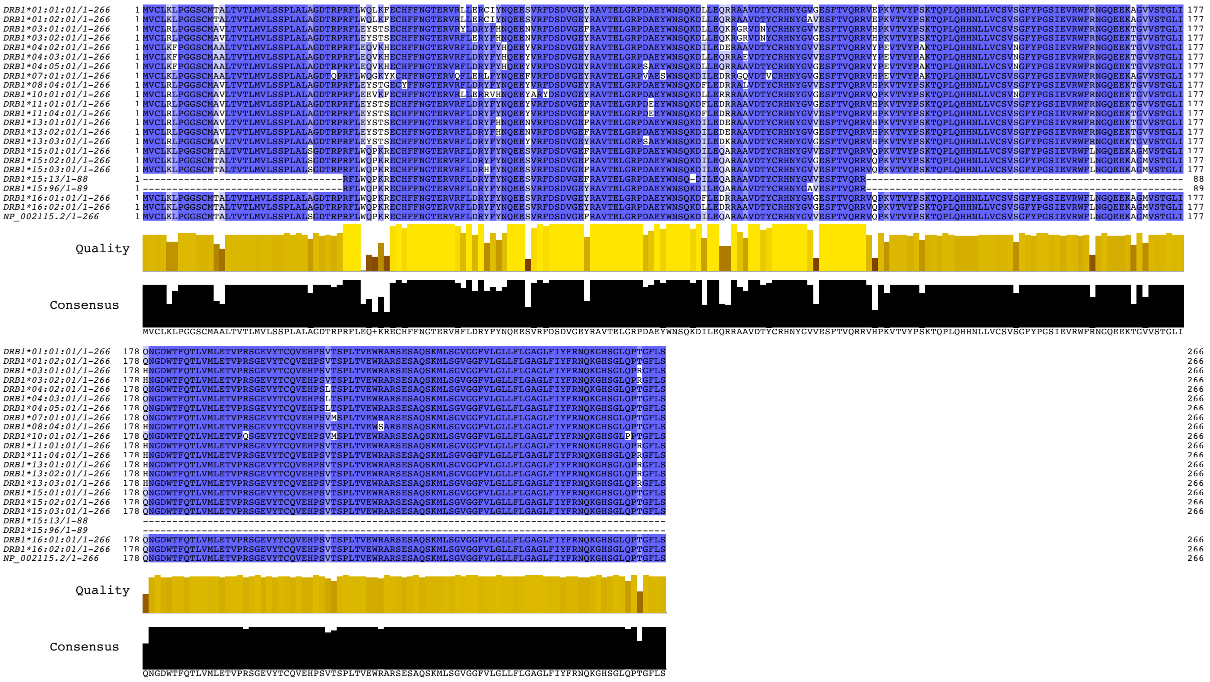


**B**

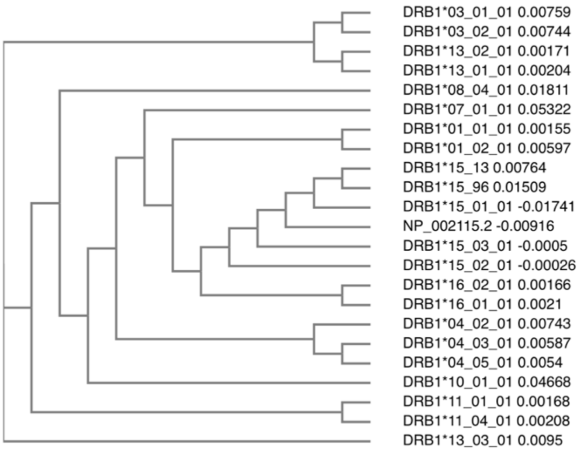


**Supplementary Figure S1. Multiple sequence alignment and phylogenetic tree of *HLA-DRB1* gene products. A,** Alignment of all HLA-DRB1 alleles analyzed in this study with the reference sequence (NP_002115.2). The color in **(A)** highlights percentage identity. **B,** Cladogram of the sequences shown in **(A)**, with the real branch length indicated by the number shown next to each allele. Sequences were retrieved from the IPD-IMGT/HLA database (https://www.ebi.ac.uk/ipd/imgt/hla/allele.html).

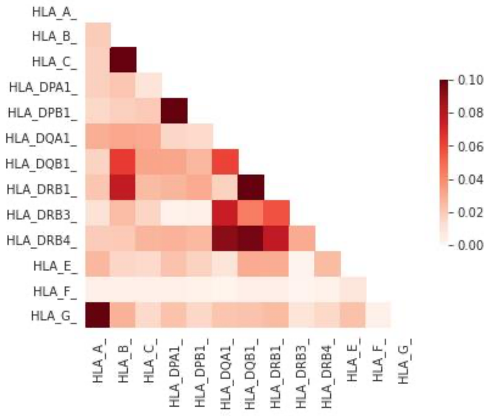


**Supplementary Figure S2.** Entropy-based linkage disequilibrium index (eLD) between the classical HLA loci in the study cohort (n = 800).

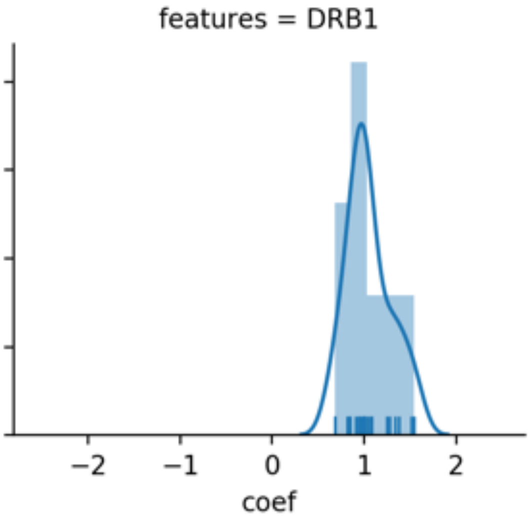


**Supplementary Figure S3. Associations between HLA DRB1 gene zygosity and the antimicrobial antibody repertoire breadth.** Distribution of the coefficients of association for HLA-DRB1 zygosity and the antibody repertoire breadth after bootstrapping (100-fold).

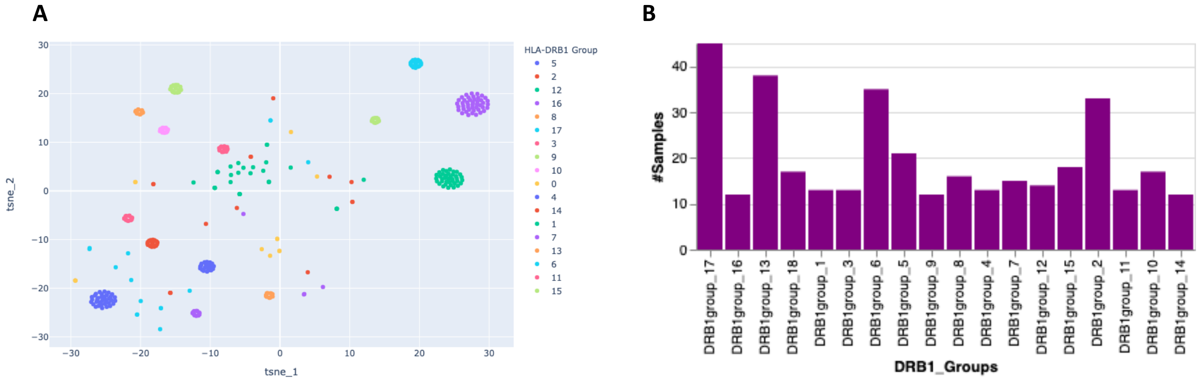


**Supplementary Figure S4. Unsupervised clustering analysis of the study cohort (n = 800) based on HLA-DRB1 genotypes using HDBSCAN. A,** tSNE projections of the samples (n = 357) with a clustering probability of 1, along with the cluster density. **B,** Bar plot depicting the sample size of each cluster (i.e., HLA DRB1 genotype group).
